# GLA-3 Mediates Stress Response in *Caenorhabditis elegans* Germ Cells: A key role of the Tristetraprolin (TTP) Family

**DOI:** 10.1101/2024.10.01.616206

**Authors:** Laura Silvia Salinas, Angel Armando Dámazo-Hernández, Enrique Morales-Oliva, Laura Ivón Láscarez-Lagunas, Rosa Estela Navarro

## Abstract

Tristetraprolin or TTP is an RNA-binding protein that possesses two CCCH-like zinc-finger domains that bind AU-rich elements to promote their degradation. One of its targets is the mRNA of tumor necrosis factor alpha (TNF-α). When TTP is absence, the TNF-α factor accumulates causing severe, generalized inflammation in knockout mice. TTP is also considered a tumor suppressor protein because regulates the expression of several mRNAs that encode for proteins involve in cell cycle regulation and it is downregulated in various types of human cancers. Under stress, TTP associates with stress granules (SGs), dynamic cytoplasmic condensates formed by liquid-liquid phase separation (LLPS) that protect mRNAs from harmful conditions. Despite TTP’s important role in mRNA turnover, much remains to explore about its participation in stress resistance in life animals that is why, we explored the role of GLA-3, one of the TTP’s homolog, in the nematode *Caenorhabditis elegans*. Nematodes lacking *gla-3*/TTP exhibit phenotypes such as progressive loss of motility, reduced brood size, and increased embryonic lethality. As well as defects in meiotic progression, and increased germ-cell apoptosis. Here we showed that the GFP::GLA-3 reporter is expressed mainly in the *C. elegans* germline, where associates with different condensates like germ granules, processing bodies, and stress granules suggesting that, like TTP, GLA-3 plays an important role in mRNA regulation in the *C. elegans* germline. Furthermore we demonstrated that GLA-3 is important for stress granules’ and processing bodies’ formation. We also show that oogenic germ cells of GLA-3 mutant animals that were exposed to heat shock resulted embryos that did not survive showing that GLA-3 plays an important role protecting germ cells from this condition. Our results demonstrate that the role of GLA-3 is conserved in *C. elegans*, and this model can be very useful for further investigating the role of this protein on the future.

## Introduction

Germ cells transmit essential information for the next generation by providing maternal mRNA and RNA binding proteins that regulate mRNA expression during early embryogenesis. Consequently, the control of translation is key in germline development and function. RNA granules or ribonucleoprotein complexes (RNPc) are biomolecular condensates formed by liquid-liquid phase separation (LLPS) that control mRNA regulation (1) (2) (3). Stress granules (SGs) are among some of the best-studied biomolecular condensates; they are assembled mainly during the arrest of translation initiation triggered by stressful conditions (4). RNA granule formation is orchestrated by proteins with intrinsically disordered regions (IDR) and/or low-complexity domains (LCD) (5). Among the key proteins that trigger SGs formation we find the RNA binding proteins, such as the T-cell-restricted intracellular antigen protein (TIA-1/TIAR) (6) and Tristetraprolin (TTP) (7), which possess prion-like domains and IDR domains, respectively, and are important for RNA granules nucleation (8) (9) (10).

The TTP family of proteins plays an important role in mRNA regulation. TTP is part of a family of CCCH tandem zinc finger proteins (TIS11) that interacts directly with the AU-rich elements (AREs) of mRNA 3’UTR (11) to promote deadenylation and, eventually, its degradation (12). For example, TTP promotes the degradation of tumor necrosis factor alpha (TNF-α) and other cytokines mRNA, explaining why mutant mice lacking TTP have severe, generalized inflammation (11). The lower expression of TTP has been also related to cancer; thus, this protein is considered a tumor suppressor in many types of cancer (13).

Under non-stressful conditions, TTP is diffusely distributed in the cytoplasm where it localizes with DCP1 (mRNA-decapping enzyme 1A) in P bodies and in the nucleus (7) (14). P bodies (PBs) are RNP complexes that contain components of the mRNA decay machinery that are present under normal growth conditions and under stress. PBs increase in size and number by fusing with other PBs or even with SGs (14) (15).

Under stress conditions, such as exposure to CCCP, nuclear TTP translocates to the cytoplasm to associate with SGs, where it colocalizes with the SGs marker TIA1 (7). TTP assembly into SGs is regulated by post-translational modifications, such as phosphorylation; for example, TTP is phosphorylated in its S52 and S178 residues to form a TTP:14-3-3 complex, which excludes TTP from SGs and inhibits the degradation of ARE-containing transcripts (7). Diverse kinases phosphorylate TTP, such as MAPKAP kinase-2 (MK2) (7), c-Jun N-terminal kinase (JNK), p38 MAP kinase, and p42 mitogen-activated protein kinase (ERK2) (16).

*C. elegans* have several genes that encode for proteins with zinc finger domains similar to TTP such as *pie-1*, *pos-1*, *mex-1*, *mex-5*, *mex-6*, and *oma-1/-2* (17) (18) (19) (20). In this work, we studied one of the TTP nematode’s homologs, GLA-3, in the *C. elegans* gonad during stress. The *C. elegans* gonad is an excellent model for studying biomolecular condensates *in vivo* because its vast size and transparency, among other features (21) (22). Previous studies identified SGs assembly in the *C. elegans* gonad during heat shock, starvation, prolonged meiotic arrest, among other conditions (23) (24) (25) (26).

By alternative splicing, *gla-3* produces three isoforms, which are expressed in the soma and germline during early embryogenesis, L4 larvae, and adult stage (27). *C. elegans* GLA-3 is a novel component of the MAP kinase MPK-1 signaling pathway required for germ-cell survival. *gla-3* loss of function or silencing exert many effects on the nematode. Among the later are found progressive loss of motility due to protein degradation in muscle, fertility issues due to defects in meiotic progression, high levels of germ cell apoptosis, and a less severe effect on embryonic lethality (27) (28) (29). Two hybrid and immunoprecipitation assays identified an association between GLA-3 and the MAP kinase MPK-1/ERK that is is required for pachytene exit during meiosis (27). Germ cells in the pachytene region of *gla-3*-mutant animals present a delay in their progression that could be caused by a misregulation of the MAPK signal (27) (29). Despite that the GLA-3 function has been studied in *C. elegans*, we do not yet know its role in the stress response in this or other organisms. Previously, we found that when we exposed young adult hermaphrodites at up to 6 h of starvation (bacterial deprivation) or 3 h of heat shock (31°C), germ-cell apoptosis increased, and ribonucleoprotein (RNP) complexes or biomolecular ocndensates, similar to SGs, were formed in the gonad (25). We evaluated the role of GLA-3 in germ cells exposed to stress, and we found that *gla-3* mutant-animals were unable to form SGs in their gonads during heat shock or starvation, suggesting that this protein is important for the formation of these condensates. Furthermore, we demonstrated that GLA-3 plays an essential role in protecting germ cells during heat shock. Using the transgene *<u>gla-</u> 3a*(tn1734[gfp::3xflag::*gla-3a*]) (DG4230) (30), we observed that GFP::GLA-3 associates with SGs during heat shock and starvation. Additionally, we observed that silencing of *mpk-1*/ERK affected the formation of SGs, suggesting that this protein might be important in regulating SG formation through GLA-3.

## Materials and Methods

### Strains

*C. elegans* strains were maintained at 20°C on NGM-Lite and fed with the *E. coli* strain OP50-1 (31). The following strains were used: wild-type variety Bristol N2, WS2974 *gla-3(ep312)* (27), and DG4230 *gla-3a(tn1734[gfp::3xflag::gla-3a])* (32).

### RNA interference

To silence the *mpk-1* gene, we obtained the clone from the RNAi library (OpenBiosystems) and the plasmid pPD129.36 was used as control (EP, empty plasmid) (33). We used the *Escherichia coli* strain HT115(DE3) to feed animals for RNAi experiments. To induce the production of double-stranded RNA we followed standard procedures (34). Briefly, bacteria cultures were grown overnight in LB broth containing 50 µg/ml of ampicillin and 12.5 µg/ml of tetracycline. To induce double-stranded RNA formation, a drop of overnight cultures was seeded onto 60 mm NGM plates supplemented with ampicillin (50 µg/ml), tetracycline (12.5 µg/ml) and IPTG (1 mM). Plates were cultured overnight at room temperature to allow the synthesis of double stranded RNA. L4 larvae were place onto NGM-lite plates containing induced bacteria and incubated at 20°C for 24 h for RNA silence.

### Stress conditions

Synchronized L1 animals were grown at 20°C on NGM-lite plates seeded with indicated bacteria until they were one-day-old adults. Then the population was separated into stressed and control groups. For starvation conditions, 1-day-old animals were transferred to NGM-lite plates without bacteria and incubated for 6 h at 20°C. For the control group, animals were kept on NGM-lite plates seeded with indicated bacteria at 20°C. For starvation recovery experiments, animals were transferred into NGM-lite plates seeded with indicated bacteria and kept at 20°C for desired hours after stress. For heat shock, one-day-old animals were transferred onto seeded plates, which were then sealed with Parafilm, and placed in a controlled temperature water bath at 31°C for 3 hours. The control (no stress) group plates were kept on seeded plates in the incubator at 20°C. For heat shock recovery experiments, plates were transferred to an incubator and kept at 20°C for indicated hours after stress. After every treatment or recovery time, animals were anesthetized with 10 mM tetramisole, mounted on 2% agarose pads and observed under an epifluorescence Nikon E600 microscope equipped with an AxioCam MRc camera or a confocal Zeiss LSM8001 microscope. The percentage of animals that presented stress granules was determined until reaching 100% of animals that form these structures.

### Immunostaining

To visualize stress granules using CGH-1 as a marker, we perfomed immunostaing as previously reported by (35). Briefly, the gonads of one-day-old animals were dissected, freeze-cracked and fixed in cold methanol (-20°C) for 1 min. Samples were fixed in a solition containing 3.7% paraformaldehyde, 80 mM HEPES, 1.6 mM MgSO4, and 0.8 mM EGTA dissolved in 1X PBS for 15 min at room temperature. After fixation, samples were washed twice with PBT, and were then blocked in PBT containing 30% normal goat serum (NGS; Sigma-Aldrich, St. Louis, MO) for 30 min. Primary antibody were diluted in PBT with NGS and the incubation was performed overnight at 4°C with rabbit anti-CGH-1 (1:1000 (36), and mouse anti-GFP (1:5000; A11120 from Molecular probes, Eugene, OR). Secondary antibody incubations were performed for 2hr at room temperature with Alexa Fluor 594-conjugated anti-rabbit IgG and Alexa Fluor 488-conjugated anti-mouse IgG (1:100; H+L, Molecular Probes, Eugene, OR). To detect DNA 1ng/µl 4′6′-diamidino-2-phenylindole (DAPI) was used. Vectashield Mounting Medium (Vector Laboratories, Burlingame, CA) was added to avoid photo bleaching before sealing the sample. At least two independent experiments were conducted with n≥50 for each condition and time point. The average percentage of animals with visible granules is depicted in the graphs.

### Quantitation of embryonic lethality after stress

Experiments were performed as previously published by (25). Hermaphrodites were grown at 20°C. They were cloned individually to plates at the mid-L4 stage. 18 to 20 hours later the young adult hermaphrodites were transferred to seeded plates which were then sealed with Parafilm and put into a controlled temperature water bath at 31°C for 3 hr. Control groups (no stress) were kept on NGM-lite plates seeded with OP50-1 at 20°C. Immediately after the stress, animals were mounted without any anesthetic onto 2% agarose pads with M9 and observed under the microscope. The embryos in the uterus and fully-grown oocytes (–1 to –3) in each gonad arm of every hermaphrodite were counted. Then, animals were recovered on NGM-lite seeded plates at 20°C and allowed to lay as many embryos as counted earlier, constituting group I. Afterward, the hermaphrodites from both strains were transferred to new plates and allowed to lay embryos for 12 hours, constituting group II. Once again, the hermaphrodites were transferred to new plates and allowed to lay embryos for another 24 hours, constituting group III. The embryonic lethality was determined as the percentage of embryos that did not hatch after 24 hours of being laid. In parallel, the embryonic lethality of hermaphrodites that were not heat-shocked was scored as a non-stress control.

### Data availability

Strains and materials will be provided upon request.

## Results

### GFP::GLA-3A/C are Expressed in the *C. elegans* Germline

To study the expression of GLA-3 *in vivo*, we used the DG4230 strain previously generated by CRISPR-Cas9 genome editing by the Greenstein Laboratory (32). This strain expresses *gla-3* isoform a and possible c, since their sequences differ only in 3 nucleotides in exon 2, fused to a *gfp* reporter in the amino-terminal and a 3xflag tag; *gfp::3xflag::gla-3a* (https://wormbase.org/species/c_elegans/gene/WBGene00011376#0-9f-10). From this point on, we will refer to this fusion protein as *gfp::gla-3* (Fig 1A). Under normal growth conditions, we observed that GFP::GLA-3 expression is restricted to the distal gonad in adult hermaphrodite animals (Fig 1B). *gfp::gla-3* is observed in the germline from the early L1 larval stage and continuing through adulthood (Fig 2A-D). GFP::GLA-3 is mainly observed in germ-cell cytoplasm and in perinuclear foci that resemble P granules (arrows in insets Fig 2 Á-D′). In the adult hermaphrodite gonad, GFP::GLA-3 expression is restricted to the distal gonad tip prior to the bend region of the gonad (Fig 2D and 3A).

**Fig 1.**
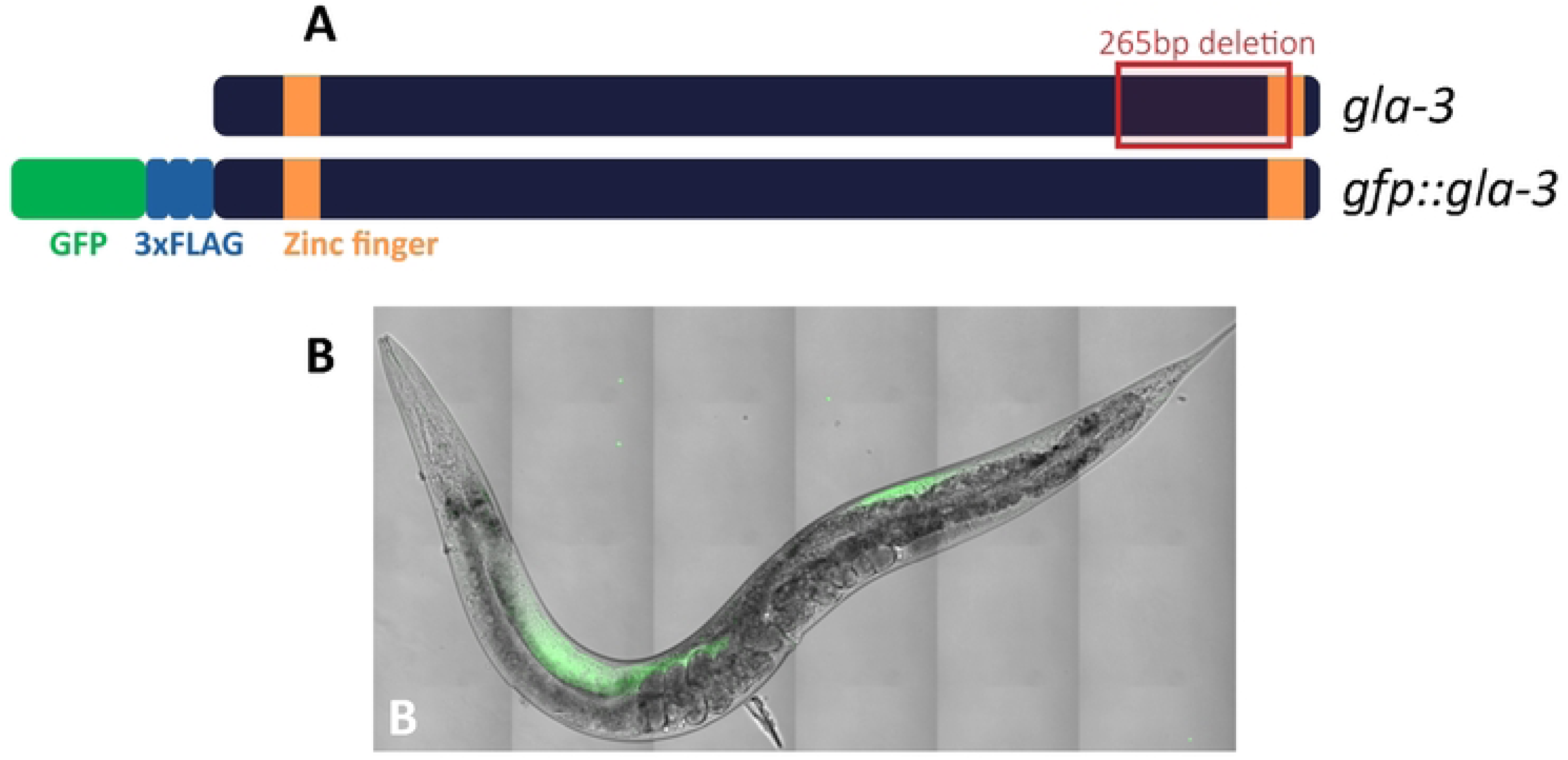
GFP::GLA-3 expression in the adult hermaphrodite. A) Scheme of the GLA-3 protein showing two zinc finger domains (orange boxes), the site of the *gla-3(ep312)* mutant deletion (outlined in red) and the GFP and FLAG tags in the GFP::GLA-3 transgene. The mutation was generated by EMS, which deleted a 265 bp region and introduced a frameshift at the new junction, affecting the three isoforms and removing part of the last two exons of *gla-3* (27). The *gfp::gla-3* construct was generated using CRISPR-Cas9 by the Greenstain Laboratory (32). The *gfp* is in the N-terminus of the *gla-3* gene (green) and carries a 3xFLAG at the N-terminus (blue). B) Merge image of a live adult animal observed in Nomarski and epifluorescence microscopy showing GFP::GLA-3 expression under normal growth conditions.

**Fig 2.**
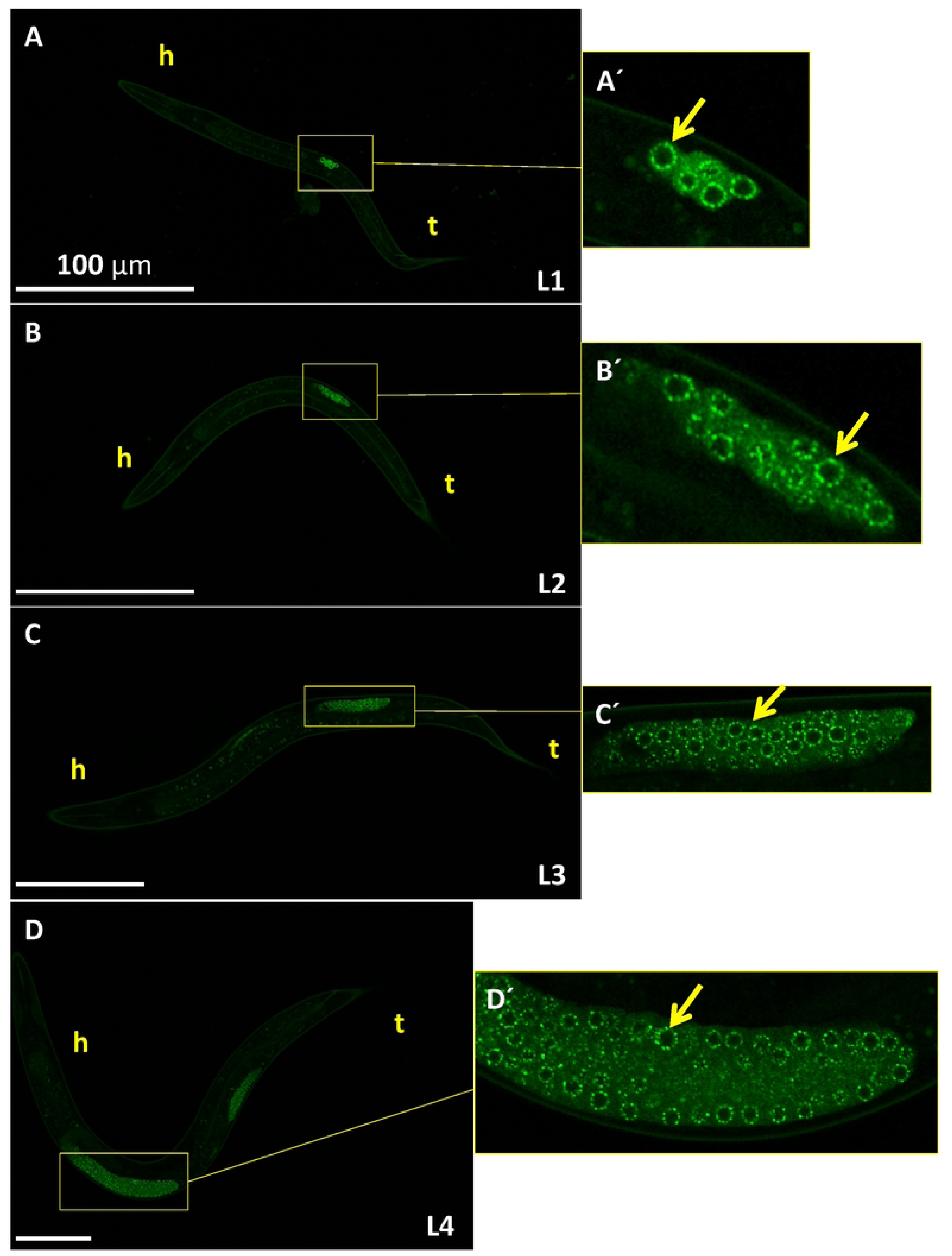
GFP::GLA-3 expression is restricted to the germline. A-D) Live animals, expressing a GFP::GLA-3 transgene at the indicated larval stages, were anesthetized and observed under confocal microscopy. Á-D′) Details of each gonad are shown at the right (yellow boxes). Arrows point toward germ cells’ perinuclear foci. h=head and t= tail. Scale bar= 100 µm.

**Fig 3.**
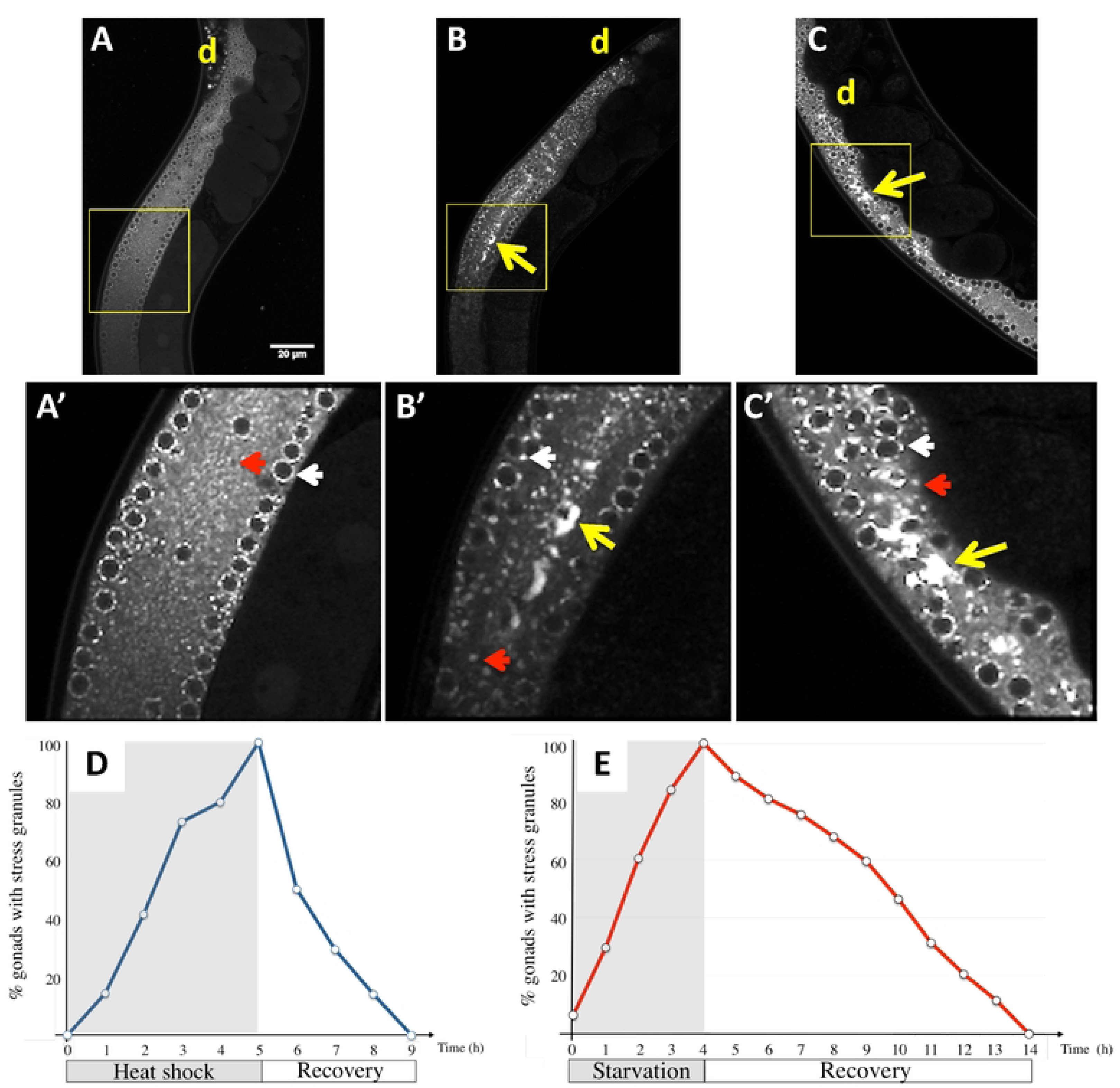
The formation of GFP::GLA-3 condensates during heat shock or starvation is transitory. (A-C) Confocal images of GFP::GLA-3 transgenic animals under control (A), heat shock (3 h 31°C) (B) or 4 h of bacterial deprivation (C). Details of each image are shown on A’, B’ and C’, respectively (yellow boxes). White arrowheads point toward perinuclear foci (P granules), red arrowheads point toward scattered cytoplasmic granules (putative storage bodies or P bodies) and yellow arrows point toward gonad core stress granules. d= distal. Scale bar= 20 µm. (D and E) *gfp::gla-3* 1-day-old hermaphrodites were exposed to heat shock (5 h at 31°C, D) or starvation (4 h with no bacteria, E). Animals were observed under the epifluorescense microscope every hour and were scored for the presence of GFP::GLA-3 gonad core granules during stress exposure and recovery, until granules were no longer observed. Two independent experiments were conducted under each condition and time point (n=50). The average percentage of animals with visible granules is depicted in the graphs.

### GFP::GLA-3 Associates with Stress Granules

To test how GFP::GLA-3 expression is affected by stress, we subjected 1-day-old *gfp::gla-3* animals to 31°C for 3 h (heat shock), 6 h of starvation (no bacteria) or kept them under control conditions (21°C with food *at libitum*). Control, heat-shocked, and starved animals were mounted for observation under a confocal microscopy. Control animals showed the expression of GFP::GLA-3 in the distal gonad in the germ cell’s cytoplasm and in perinuclear granules similar to P granules (white arrowhead Fig 3A and A′). We also observed GFP::GLA-3 expression in germ cell’s cytoplasm as a scattered pattern (red arrowhead Fig 3A and A′) that could be putative storage bodies or P bodies (35) (36). Heat-shocked animals exhibited larger and fewer perinuclear granules (white arrowhead), larger scattered granules (red arrowhead), and large *gfp::gla-3* granules in the middle of the gonad core (yellow arrow) (Fig 3B and B′). Starved animals showed perinuclear granules (white arrowhead), scattered punctuated granules in the gonad core (red arrowhead), and large granules in the middle of the gonad core (yellow arrow) that from this point on we will refer to as stress granules (SGs) (Fig 3C and C′).

A feature of stress granules is that they are transitory (6) (25); therefore, we examined their assembling and disassembling kinetics. For this, GFP::GLA-3 animals were subjected to heat shock or starvation conditions, as described previously, until all animals have formed stress granules, 5 h for heat shock and 4 h for starvation, respectively (Fig 3D and E). Animals were mounted every hour under the microscope to quantify the percentage of gonads that showed SGs formation. Stress granules, formed under heat-shock conditions, disassemble gradually in 4 h; in contrast to those formed under starvation, which lasted for 10 h. Our data showed that the association of GFP::GLA-3 to gonad core stress granules under stress conditions is reversible.

To test whether GFP::GLA-3 associates with stress granules in the gonad during stress conditions, we used the RNA helicase CGH-1 as a marker of stress-granules as previously described (25). Gonads of 1-day-old GFP::GLA-3 animals subjected to stress were dissected, fixed, and co-stained with CGH-1, GFP antibodies and DAPI. We observed that CGH-1 and GFP::GLA-3 colocalized in gonad core stress granules during heat shock and starvation (Fig 4 A-C’).

**Fig 4.**
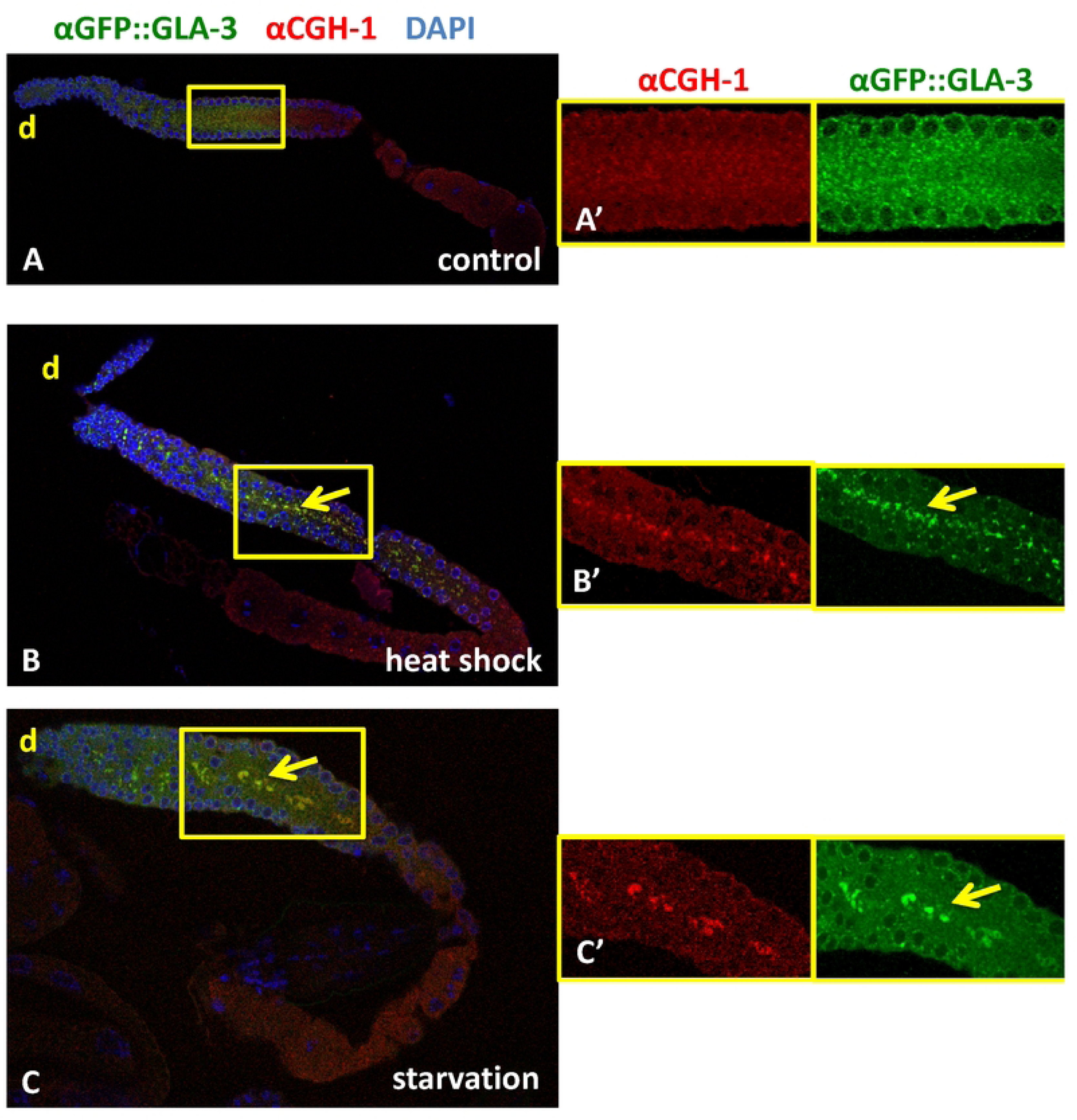
GFP::GLA-3 colocalizes with CGH-1 in stress granules. 1-day-old animals expressing a GFP::GLA-3 fusion protein were subjected to control conditions (21°C), heat shock (31°C for 3 h and 1 h recovery), or 6 h of starvation (no bacteria). After treatment, the gonads were dissected, fixed and co-stained with CGH-1 and GFP antibodies. A-C) Merge confocal images of dissected gonads under the described conditions. Details of each channel (yellow boxes) are shown in A’-C’. Yellow arrows point toward colocalization of CGH-1 and GFP::GLA-3 expression in stress granules.

### GLA-3 is Required for Gonad Core Stress Granules Formation During Stress Conditions

TTP, the GLA-3 homolog in mammals, is involved in stress-granules assembly (7). We found that GLA-3 is also required for stress-granule formation during heat shock and starvation in the *C. elegans* gonad. To carry out these experiments, we used an antibody against CGH-1 as a stress-granule marker. Under control conditions, CGH-1 is found in the germ-cell cytoplasm in a punctated pattern known as storage bodies or P bodies (35) (36) (Fig 5A, A’ and 6A and A’). Under stress conditions, CGH-1 accumulates in stress granules in the gonad core and the oocytes (Fig 5C, C’ and Fig 6C and 6C’) (25). N2 and *gla-3(op312)* 1-day-old animals were subjected to control conditions (21°C), heat-shocked (3 h at 31°C), or starved for 6 h (no bacteria). After stress, the gonads of control and treated animals were dissected, fixed and stained with a rabbit anti-CGH-1 antibody and DAPI to detect DNA (36). In heat-shocked wild-type animals, CGH-1 accumulated in SGs in the gonad core (70%) and oocytes (81%) of the gonads observed (Fig 5C and C’). In contrast, all of the gonads from *gla-3(op312)* animals did not form SGs under control or heat-shock conditions (Fig 5B, B, D and D’). A small percentage of *gla-3(op312)* gonads revealed CGH-1 granules in oocytes under control conditions (12%), while the majority demonstrated SGs in the oocytes under heat shock (94%) (Fig 5D and D′). We noticed that the small condensates known as storage granules or P bodies are diminished in number in *gla-3* animals under both control and heat shock conditions (Fig 5A′, B′, ĆAnd D’); suggesting that GLA-3 is important for CGH-1 association to P bodies and SGs during heat shock.

**Fig 5.**
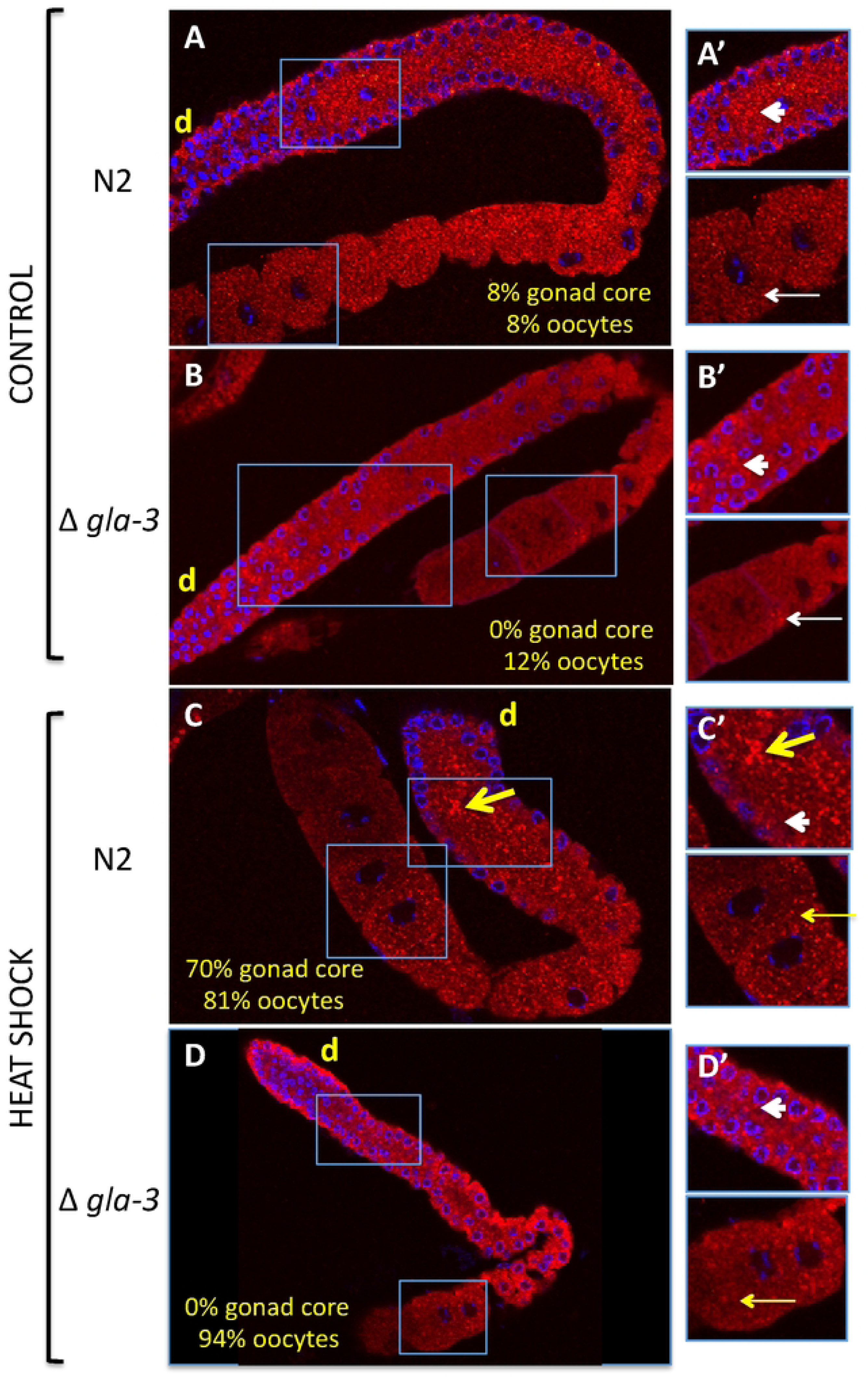
*gla-3* mutant animals are unable to form gonad core stress granules under heat shock. **A-D)** Wild-type (N2) (A and C) and *gla-3(op312)* (B and D) 1-day-old animals were maintained under control conditions (20°C) (A and B) or exposed for 3 h to 31°C to induce heat shock (C and D). After heat shock, the gonads were extruded, fixed, and stained with a rabbit anti-CGH antibody (red) and DAPI (blue). Samples were mounted under the confocal microscopy for observation and quantification of animals showing stress granules. The percentage of gonads that presented CGH-1 stress granules in the gonad core or oocytes is indicated in each panel. A’-D’) Details of each gonad are shown at the right for each picture (blue boxes). Yellow arrows point toward CGH-1 stress granules in the gonad core (thick yellow arrows) or oocytes (thin yellow arrows). White arrowheads (gonad core) and thin white arrows (oocytes) point toward condensates present in control conditions (storage bodies). d=distal.

**Fig 6.**
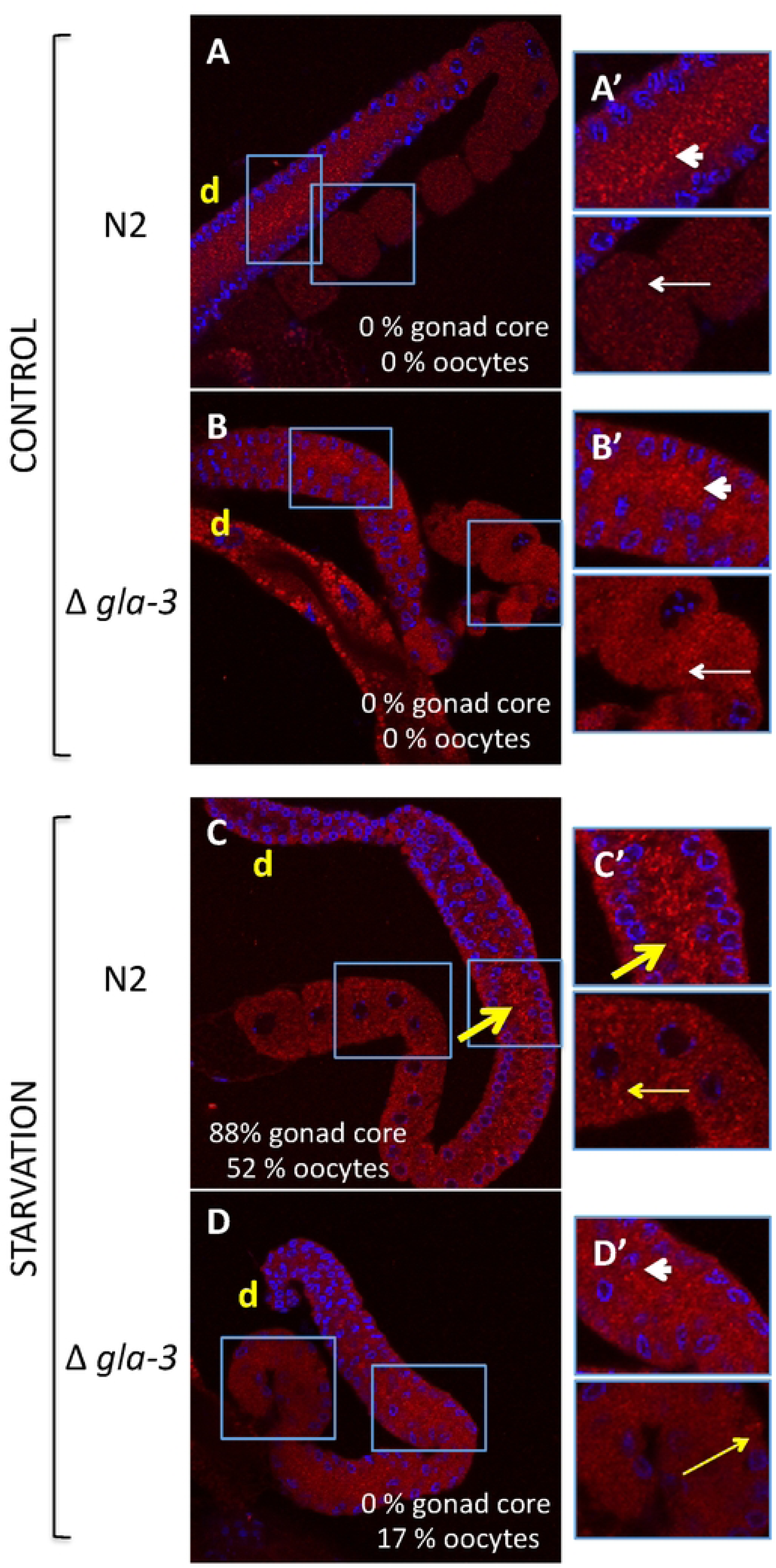
*gla-3* mutant animals do not form gonad core CGH-1 stress granules under starvation. **A-D)** Wild-type (N2) (A and C) and *gla-3(op312)* (B and D) 1-day-old animals were maintained under control conditions (20°C) (A and B) or were starved for 6 h (B and D). After starvation, the gonads were extruded, fixed, and stained with a rabbit anti-CGH antibody (red) and DAPI (blue). Samples were mounted under confocal microscopy for observation and quantification of animals showing stress granules. The percentage of gonads that presented CGH-1 stress granules in the gonad core or oocytes is indicated in each panel. Details of each gonad are shown at the right of each picture. A’-D’) Details of each gonad region are depicted to the right of each picture. Yellow arrows point toward CGH-1 stress granules in the gonad core (thick arrow) or oocytes (thin arrows). White arrowheads (gonad core) and thin white arrows (oocytes) point toward condensates present in control conditions (storage bodies). d=distal.

For the starvation experiments, gonads of wild type and *gla-3(op312)* animals under control conditions exhibited comparable CGH-1 expression to those of the wild-type animals (Fig 6A-B’). After starvation, wild-type animals showed SGs in the gonad core (88%) and oocytes (52%) (Fig 6C and C’). In contrast, gonads from *gla-3(op312)* animals did not demonstrate SGs in the gonad core, whereas only 17% showed large CGH-1 granules in oocytes (Fig 6D and D′). Similar to heat-shocked animals, the gonads of *gla-3* mutant animals also showed fewer P bodies. Collectively, our data demonstrated that GLA-3 is necessary to form SGs and P bodies under heat shock and starvation.

### SGs Do Not Assemble Efficiently in *mpk-1* Knockdown Animals

To study whether the MAPK MPK-1, *C. elegans* ERK homolog participates in stress-granule formation, we silenced *mpk-1* in *gfp::gla-3* hermaphrodites. One-day-old control (EV) and *mpk-1(RNAi)* animals were maintained at 20° C or heat-shocked for 3 h at 31° C. After treatment, the gonads were extruded in M9 buffer containing DAPI. Samples were mounted to quantify the formation of SGs. At 20°C, the vast majority of control (EV) or *mpk-1(RNAi)* animals did not form SGs (Fig 7A, B, E). Most of heat-shocked control *gfp::gla-3* animals exhibited well-formed SGs (77%); in contrast only 30.2% *mpk-1(RNAi)* animals showed condensed SGs while 69.8% accumulated GFP::GLA-3 expression at the center of the gonad, but no visible well-defined SGs were observed in their gonads (dispersed granules) (Fig 7C, D, E). Our data suggest that MPK-1 is important for the formation of well-defined GFP::GLA-3 SGs during heat shock.

**Fig 7.**
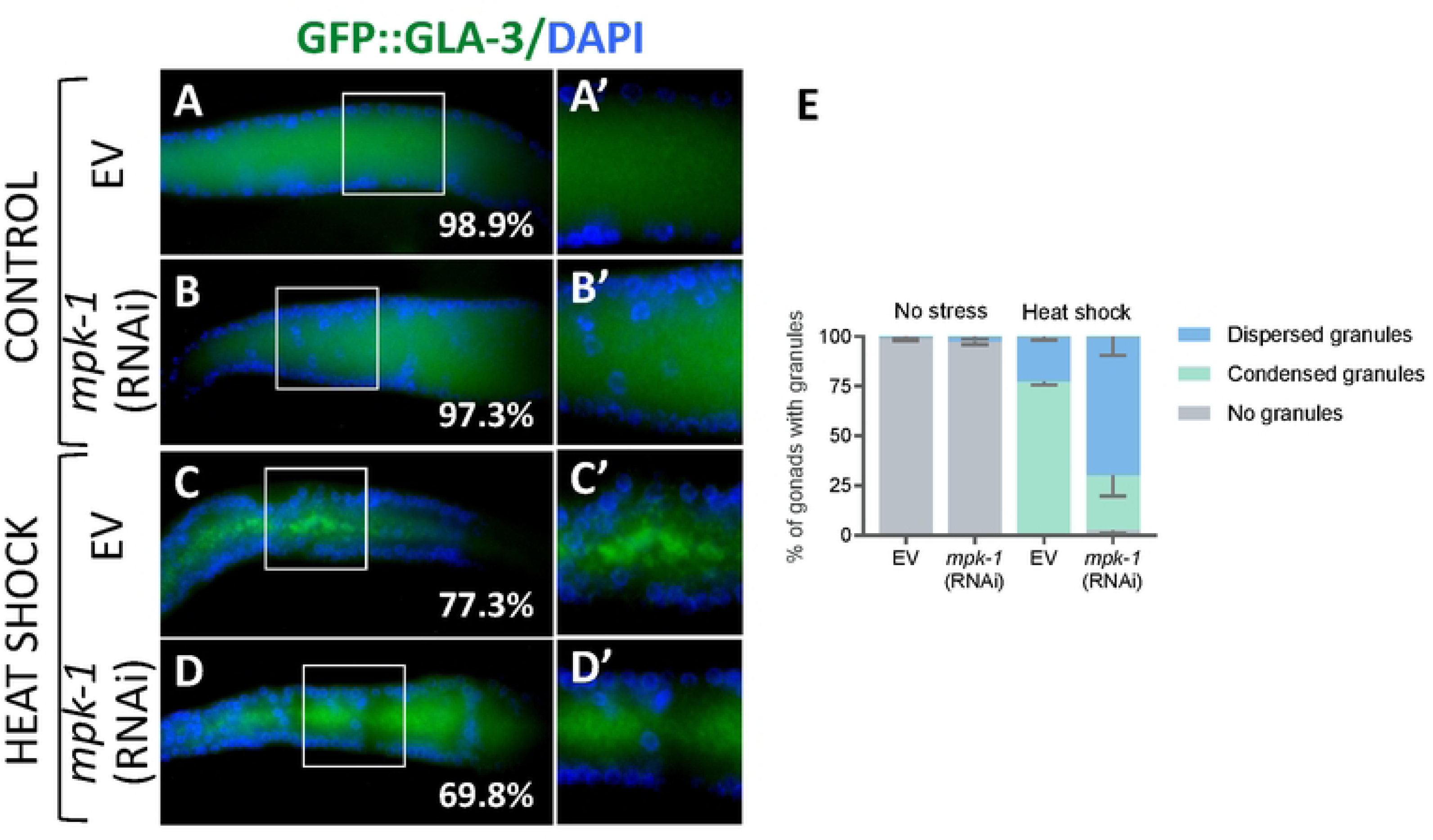
The gonads of *mpk-1(RNAi)* showed fewer GFP::GLA-3 gonad-core stress granules under heat shock. GFP::GLA-3 animals were fed for 24 h bacteria expressing *mpk-1* dsRNA or an empty plasmid since the L4 larval stage to induce RNA interference. One-day old animas were maintained at 20°C as control or were exposed to 31°C for 3 h. After treatments, the gonads were extruded in M9 buffer with DAPI to visualize nuclei. Samples were mounted under the epifluorescence microscope to quantify stress-granules formation. A-D Merge images of extruded gonads showing GFP::GLA-3 expression and DAPI for each condition. A’-D’ A detail of each gonad is depicted at the right (white boxes). The percentage of gonads showing a phenotype is shown for each panel. E) Graph showing the percentage of gonads that formed condensed, dispersed, or no granules for each genotype is shown for each condition.

### GLA-3 Protects Germ Cells from Stress

To study the role of GLA-3 in the germ cells stress response in *C. elegans*, we subjected 1-day-old adults to heat shock and followed animals progeny over several hours afterwards to quantify embryonic lethality as described by (36). After heat shock, the animals’ progeny was followed for 48 hr (see Materials and Methods). Wild-type animals showed high embryonic lethality in the first group and much lower in the group II, while the group III exhibited nearly no embryonic lethality (Fig 8). In contrast, *gla-3(op321)* animals continued to show high levels of embryo lethality even several hours after the heat shock was performed in groups II and III (Fig 8). These data suggest that GLA-3 is important for protecting germ cells from heat shock.

**Fig 8.**
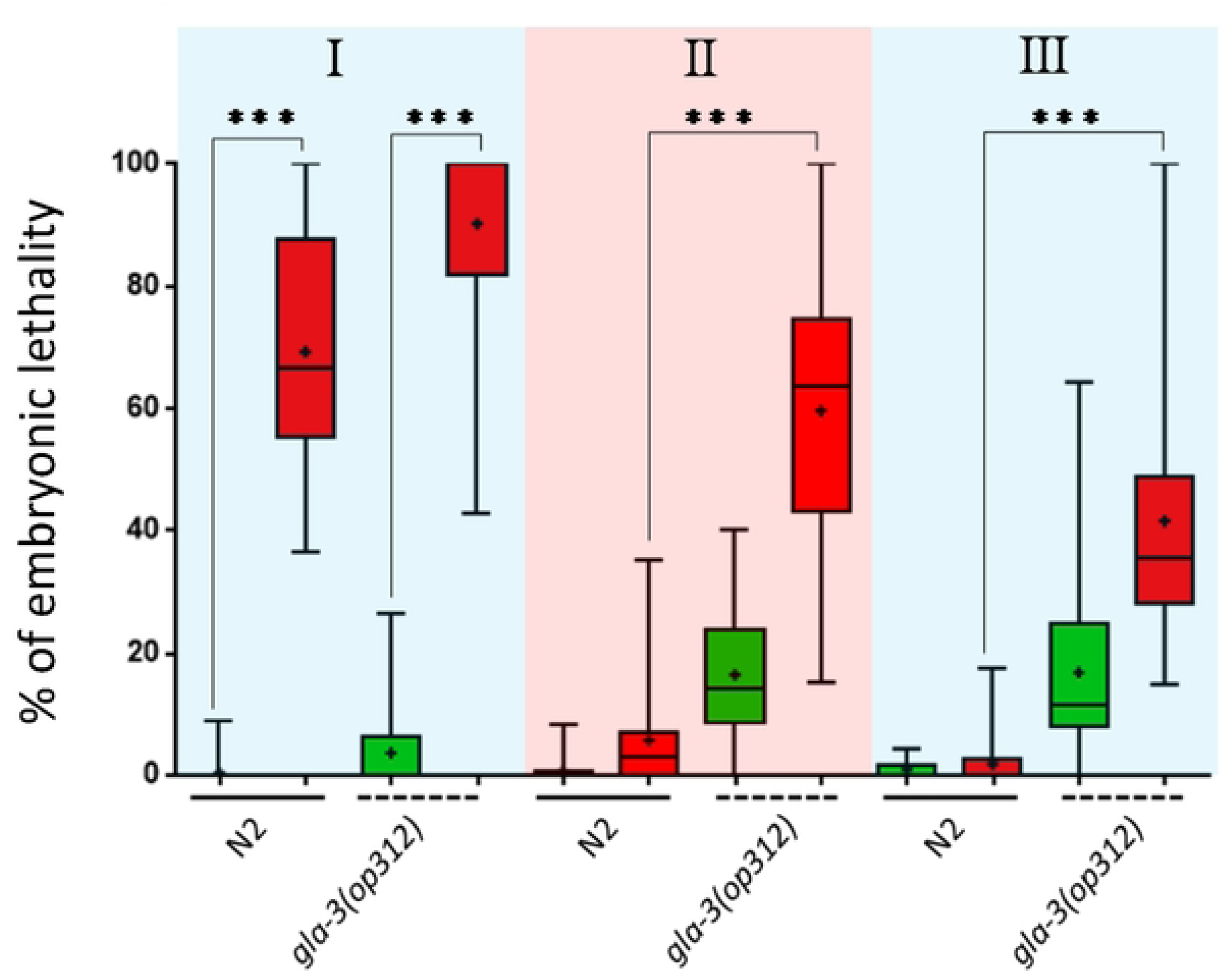
GLA-3 protects germ cells from heat shock. One-day-old N2 and *gla-3(ep312)* animals were subjected to 3 h of heat shock at 31°C or were maintained at 20°C as control condition. After stress, the animals were recovered to 20°C on seeded plates and moved to new plates every 24 hrs. Embryonic lethality was quantified after 24h of transferring the parental animals to new plates. Animals were classified in three groups, I-III, depending on the timing that has past after the heat shock. Green bars represent animals maintained at 20°C, while red bars represent animals that were exposed to heat shock. The boxes represent the interquartile range (IQR) from 25%-75%, and the bars extend from the minimal to the maximal value. The three asterisks indicate significance with a value of p<0.05. The data shown are the result of three independent experiments. The statistical model employed was one-way ANOVA with the Bonferroni multiple comparisons test.

## Discussion

The TTP family of proteins has been extensively studied *in vitro*; however, its role in the germline and under stress conditions remains poorly understood. Here we show that the TTP homolog in *C. elegans*, GLA-3, plays an important role in germ cells. Germ cells from GLA-3 mutant animals are more sensitive to heat shock and are unable to form stress granules. We also show that, similarly to its mammalian homolog, GLA-3 associates with different condensates in the germline, suggesting that GLA-3 conserves its role in *C. elegans* as an important regulator of mRNA expression.

### GLA-3 Associates with Stress Granules in the Germline

Germ cells have several types of biomolecular condensates that control mRNA expression during gametogenesis (22) (2). Here, we demonstrate that GLA-3 associates with P granules in germ cells, as well as with condensates that are similar to P bodies and stress granules, suggesting that like TTP, GLA-3 plays a key role in mRNA regulation in the germline.

P granules are generically known in other organisms as germ granules, which are present in the majority of germ cells. In 1998, Brangwynne et al. demonstrated that P granules shown liquid properties (1). This discovery was a landmark for the field of liquid-liquid phase separation (3). Germ granules are sites of maternal inherence, mRNA sorting, post-transcriptional regulation, and small RNA biogenesis (recently reviewed by (3)). The association of GLA-3 with germ granules opens the possibility that this protein plays an important role in any of these functions. Interestingly, the transgene GFP::GLA-3 is expressed mainly in germ cells that are undergoing pachytene; particularly in this region, transcription is very active. It is possible that GLA-3 regulates the expression of maternal mRNAs that are transcribed in this region.

*C. elegans* germ cells also posses condensates similar to P bodies that have been observed using proteins such as CGH-1, CAR-1, DCP-2, among others (35) (36) (37) (23). Here we show that GLA-3 is expressed in small granules within the germ cells that resemble P bodies. Several lines of evidence have demonstrated that TTP associates with P bodies in mammals. For example, TTP binds to ARE-containing mRNA to promote their decay and drive them to P bodies (38). Additionally, when mRNA decay is inefficient, TTP sequester ARE-mRNA in PB. Furthermore, when enzymes related to mRNA degradation, such as XRN1 or DCP2, are knockdown, ARE containing mRNA accumulated in P bodies along with TTP (39). TTP can also trigger PBs formation when cells are treated with cycloheximide, which normally disrupts PB formation due to translational arrest (39). Due to the association of GLA-3 with PB-like condensates, it is possible that GLA-3 might play similar roles to those TTP in the nematode’s germline. Whether GLA-3 promotes mRNA degradation of ARE containing mRNA like its mammalian homolog remains to be elucidated.

Upon stress, *C. elegans* germ cells trigger the formation of large granules that reveal many features of mammalian-stress granules (reviewed by (22)). For example: 1) gonad-stress granules have several conserved stress-granule markers such as TIA-1 and CGH-1 (an RNA helicase) (25) (40) (23), G3BP-1 (41) and PAB-1 (23); 2) Similar to SGs in mammals, germline SGs are formed in response to translational arrest (25); 3) they are dissembled in the presence of cycloheximide and assembled in Puromycin (25), and 4) similar to SGs, gonad-stress granules formation is transitional (25).

Intriguingly, gonad’s stress granules in *C. elegans* also have some markers of P bodies, such as DCP-2 (23), CAR-1/RAP55 (23) and CGH-1/p53 (23) (25). Here we show that GFP::GLA-3 co-localizes with CGH-1, confirming a close relationship between SG and PB in the *C. elegans’* germline. It has been shown that stress granules and PBs are closely related in yeasts and mammals (42).

Although stress granules observed in the *C. elegans* gonad core and oocytes associate with the same proteins; they diverge in some aspects. Particularly, gonad core stress granules require TIAR-1 or GLA-3 for their formation, while oocyte-stress granules do not ((25) and this work, respectively). Oocyte condensates are also observed in hermaphrodites that do not have sperm; this condition, known as arrested oogenesis, is present in old animals (more than 3-days-old) or some genetic backgrounds that feminized the germline (23). Large condensates observed in arrested oocytes showed the same markers as PB, and SGs in addition to MEX-3, an RNA binding protein important for germline and embryonic development (43) (14). However, large condensates in arrested oocytes appear to have liquid-gel consistency because they do not reconstitute after photobleaching as fast as condensates observed in the pachytene germ cells (44) (45).

### The TTP Family of Proteins plays an important Role in Stress Granules Formation

There are some proteins that participate in the condensation of granules in the *C. elegans* germline, but much remains to be learned about this mechanism. P granule nucleation is the best-characterized in *C. elegans*, and mainly requires the RNA binding proteins PGL-1, -2, and -3 and the DEAD box RNA helicases GLH-1 and GLH-4 (reviewed by (46)). PGL and GLH proteins possess intrinsically disordered or low complexity domains that bestow on P granules a liquid-like behavior (47). During late oogenesis and early embryogenesis, P granules detach from the nuclear pores and get surrounded by the intrinsically disordered proteins MEG-3 and MEG-4 conferring them a gel-like consistency that bestows them more resistant (48). Despite that MEG-3/MEG-4 are present in oocytes, surprisingly, these proteins do not localized to all large oocyte granules, and they are not necessary for PGL-1, CGH-1, or MEX-3 association to these RNP (45).

A key player in the condensation of large oocyte granules during arrest oogenesis conditions is the PUF family of translational repressors (44). The PUF proteins are necessary for the condensation of CAR-1 or the deadenylase CCF-1 into large oocyte granules. Particularly, PUF-5 contributes to the condensation of MEX-3 and MEG-3 into large granules in arrested oocytes; however, it is not important for PGL-1 condensation into these particles (45). Intriguingly, neither MEX-3, TIAR-1, nor GLA-3 play a role in oocyte SG formation in arrested oocytes or under stress conditions ((45) (25) and this study). It is possible that PUF-5 or other members of its family could play a role in oocyte SG formation observed under other stress conditions, such as starvation and heat shock.

The DEAD box RNA helicase CGH-1 plays an important role by maintaining arrested oocytes in a “liquid-gel”-like consistency, because when *cgh-1* is missing, large CAR-1 granules, present in arrested oocytes, acquired a square-sheet consistency that does not reconstitute after photobleaching, suggesting that they might have a more solid phase (44).

The role of TTP in stress-granule formation has been described in mammals. The overexpression of TTP induces stress-granule formation even in the absence of stress (7). However, the association of TTP with stress granules in mammals depends on the specific condition being tested. For instance, oxidative stress induced by FCCP localizes TTP to stress granules; in contrast, another type of oxidative stress induced by arsenite excludes TTP from these condensates (7). TTP exclusion of SG during arsenite exposure is due to the activation of the p38-MAPK/MK2 kinase cascade, which triggers TTP phosphorylation, disabling it to associate with these condensates (7). Here we show that the role of GLA-3 in stress-granule formation during stress is conserved in *C. elegans*; notwithstanding this, further studies are required to understand exactly how this family of proteins can influence the formation of these condensates.

### MPK-1/ERK Is Important for Gonad-core Stress-granule Condensation

MPK-1 is the *C. elegans* homolog of the mammalian ERK1/2 serine/threonine kinases. In *C. elegans*, *mpk-1(lf)* mutant animals are sterile because their germ cells fail to exit the pachytene stage of meiosis I (49). The germ cell exit from pachytene is also a requisite for apoptosis, and that is perhaps the reason why *mpk-1(lf)* mutant animals do not undergo physiological germ-cell apoptosis (50). Kritikou et al. elegantly showed that MPK-1 interacts both genetically and biochemically *in vitro* and *in vivo* with GLA-3 (27). MPK-1/GLA-3 interaction is important for germ-cell apoptosis, vulval development, and muscle integrity.

In an *in silico* analysis performed by Kritikou *et al*., (27) the authors revealed that GLA-3a contains a putative MPK-1 phosphorylation site at position 66-75 (LRKVVRIDR), while GLA-3b has two phosphorylation sites at positions 66-74 (LRKTKI) and 99-107 (LRKVVRIDE). We found that GLA-3a possesses a putative ERK1 phosphorylation site at position S215. It would be interesting to probe whether this phosphorylation site is important for GLA-3 function. Despite that GLA-3 interacts with MPK-1 and has putative sites of phosphorylation, GLA-3 protein accumulation does not appear to be affected in *mpk-1* pathway-deficient animals. Similarly, the lack of GLA-3 does not affect MPK-1 expression or activation (27). Here we demonstrated that *mpk-1(RNAi)* animals are deficient in GFP::GLA-3 gonad-core stress-granules formation (Fig 7 C-D). These data suggest that GLA-3 phosphorylation might trigger stress-granules assembly, although we cannot rule out that MPK-1 phosphorylates other protein or proteins that trigger stress-granules accumulation independently of GLA-3. Previously, Jud et al. showed that MPK-1 expression correlates with the formation of large RNP in arrested oocytes (23). In high MPK-1 expression, large RNP are formed in arrested oocytes, while at lower levels, these particles are not observed. Overall, it is possible that MPK-1 might be triggering the formation of RNP through multiple mechanisms.

The GLA-3 homolog in mammals, TIS11, is also phosphorylated by MAPKMAP kinase 2 (MK2), ERK2, p38, and JUNK (51) (52) (53). Phosphorylated TTP is recognized by the 14-3-3 protein, a low-molecular-weight adaptor that recognizes phosphorylated proteins and promotes changes in localization and function (7). As a consequence of this, phosphorylated TTP cannot bind the CCRR4-NOT complex, impairing the participation of TTP in mRNA turnover and affecting its localization to stress granules (53). As a result of TTP phosphorylation, its mRNA targets are stabilized (54) (55) (7). TTP phosphorylated by MK2 (p38 pathway) binds less avidly to the ARE of TNFα mRNA, thus, stabilizing its transcript (54).

What are the consequences of a deficiency in condensate formation for an organism? This is an intriguing and fascinating question that remains to be fully elucidated. However, it is not easy to answer because the majority of proteins that contribute to condensate formation play multiple roles in RNA regulation; therefore discerning between their functions inside and outside of a condensate is by no means straightforward. Nevertheless there is some light that point toward the role of condensates in the *C. elegans* germline. For example, when germ-granules nucleators such as *pgl-1* and *glh-1/-4* are absent animals exhibit sterility (46). In other cases, the consequence is subtler, such as in *meg-3* and *meg-4* mutant animals that develop a normal adult germline, but that are unable to carry on normal miRNAs biogenesis, which results in sterility over several generations (48). Silencing genes that participate in MEX-3-granule assembly results in embryonic lethality (56). Lacking TIAR-1 and GLA-3 proteins affects germ-cells quality when exposed to heat shock ((25) and this work). In an attempt to answer this question, we disrupted the prion-like domain of TIAR-1 in *C. elegans* (26). Prion-like domains are important for condensate formation (57). We found that SGs in the *C. elegans* gonad still formed, although their consistency shifted toward a more liquid state. TIAR-1 prion-domain mutant animals continue to exhibit low fertility but showed increased embryonic lethality, suggesting that the role of this protein in condensates may be critical for these functions. Understanding the role of condensates in living organisms requires the use of specific assays, which underscore the importance of developing whole-animal models for studying this phenomenon.

## Acknowledgments

We thank members of the Navarro lab for their insightful comments on the development of this project. This work was supported by grants to R.E. Navarro: Programa de Apoyo a Proyectos de Investigación e Innovación Tecnológica, Universidad Nacional Autónoma de México (PAPIIT-UNAM; IN208918 and IN210821) and Consejo Nacional de Ciencia y Tecnología (CONACyT-México; CF-101731). We thank the *Caenorhabditis* Genetics Center (CGC), which is funded by NIH-Office of Research Infrastructure Programs (P40 OD010440). We would like to express our gratitude to Keith Blackwell and L. Paulette Fernández-Cardenas for generously providing the CGH-1 antibody.

